# Flaming as part of aseptic technique increases CO_2 (g)_ and decreases pH in freshwater culture media

**DOI:** 10.1101/780353

**Authors:** Brittany N. Zepernick, Lauren E. Krausfeldt, Steven W. Wilhelm

## Abstract

Aseptic technique has historically served as a fundamental practice in microbiology, helping maintain culture purity and integrity. This technique has been widely encouraged and employed for use with cultures of heterotrophic bacteria as well as freshwater and marine algae. Yet, recent observations have suggested these approaches may bring their own influences. We observed variations in growth amongst replicate experimental cyanobacterial cultures upon flaming of the culture tube opening during sample transfer and collection. Investigation revealed the pH of culture media had decreased from the initial pH established during media preparation. Flaming of sterile culture media alone confirmed a significant decrease, by as much as 1.7 pH units, and correlated with increased flaming events over time. We hypothesized that the causative factor was the introduction of carbon dioxide into the media. To test this hypothesis, qualitative and quantitative analyses were used to identify the primary driver of pH decline. We further assessed the direct effects of flaming and subsequent pH changes on *Microcystis aeruginosa* cultures, showing flame-driven pH changes and/or the introduction of carbon dioxide influenced experimental results. Our observations provide a cautionary tale of how classic and well-accepted approaches may not always provide the results promised, suggesting new approaches may be necessary in research areas assessing pH or carbon related-effects on microbial communities.

## Introduction

Aseptic technique has served as a foundational method of microbiological research for decades (Harrigan and McCance 1966). Defined as a series of steps to prevent contamination during the manipulation of cultures and sterile culture media (Madigan 2015), aseptic technique commonly includes flame sterilization of the opening of culture vessels (Bykowski and Stevenson 2008). Flame sterilization consists of passing a culture vessel opening through a flame to prevent the introduction of microbial contaminants to a sample and is an approach that is used in a variety of laboratories, hospitals, and industrial facilities. Yet, upon review of the literature, we found a general lack of consensus within the scientific community as to the exact method by which aseptic flaming should be performed. While flaming immediately before and after sub-samplings or culture transfers is specified in a variety of publications (Andersen 2005; Madigan 2015; Sanders 2012), the exact number of times required for the mouth of the vessel to pass through a flame is generally not specified. Additionally, while some argue flaming must be performed by passing the tube through the inner blue section of the flame (Bykowski and Stevenson 2008), others instruct the tube to be passed above the flame (Coté 1998). Further variances in flaming technique arise in the recommendation of a wait time prior to re-capping the culture post-flaming (Coté 1998), while others recommend immediate recapping to minimize introduction of airborne contaminants (Harrigan and McCance 1966). In fact, the primary purpose of flaming in itself is often disputed within the literature, as some sources claim it is to combust any contaminants located on the mouth of the vessel (Coté 1998), while others indicate it is to create an upwards convection current to prevent atmospheric contaminants from entering the tube (Madigan 2015; Sanders 2012). These variances amongst techniques create a lapse in methodological consistency within the scientific community. Additionally, the discrepancies may be exacerbated by confounding factors that are unintentionally applied because of flaming. For example, the practice of flaming in aseptic technique is mandated in the handling of freshwater media and phytoplankton cultures (Andersen 2005). Yet, previous studies have demonstrated aseptic flaming may have detrimental implications on algal cultures, such as decreased growth rates and increased cell death in cyanobacterial *Prochlorococcus spp.* cultures linked to flame-generated peroxides (Morris and Zinser 2013).

In the present study, we assessed the effects of flaming as part of aseptic technique on freshwater culture media and cyanobacterial cell growth. The extent of effects were assessed as follows: (1) the effects of aseptic flaming on different freshwater growth media taken from the scientific literature, (2) the primary mechanism responsible for observed declines in pH, (3) the effects of aseptic flaming on the growth of *M. aeruginosa* cyanobacteria cultures. Based on our observations, recommendations are made in the form of modifications and alternative approaches to aseptic flaming to effectively sterilize while eliminating the potential introduction of confounding variables.

## Materials and procedures

### Freshwater media selection and preparation

Four freshwater culture media commonly used for phytoplankton culturing (known as C, CB, CT, and CSi) were selected as medium candidates: recipes were all taken from a publication of the *NIES-Collection Microbial Culture Collection* (Watanabe 1997). The components and concentrations for each media are identical except for the buffer type and concentration, allowing for direct comparisons of the specific buffers in response to flaming (**Table 1**). Briefly, the major constituents of C media include 1.5×10^−4^ g/mL Ca(NO_3_)_2_·4H_2_O, 1.0×10^−4^ g/mL KNO_3_, 5.0×10^−5^g/mL Na_2_·*β*-glycerophosphate, 4.0×10^−5^ g/mL MgSO_4_·7H_2_O, and trace metals (trace metals adapted from original protocol slightly (Steffen, 2014). To maintain comparability, silica was not included in CSi media used in this study. Beyond these media, BG-11 medium was also tested and prepared in accordance to a standard protocol (Andersen 2005) as it is another media commonly used for cyanobacteria culture work. In addition, we assessed the effects of flaming on three commonly used heterotrophic bacterial growth media: Lysogeny broth (LB) (Bertani 1951), Nutrient broth (Sambrook et al. 1989) and Minimal media (M9) (Miller 1972). All media were prepared in 1 L volumes, titrated to a pH of 8.2 with 1 M NaOH and autoclaved. After cooling to room temperature and allowing for equilibration (1 day) the pH of each medium was confirmed *via* the sub-sampling a 10 mL volume and analysis using a calibrated laboratory pH meter (Mettler Toledo Seven Compact). Vitamin additions pertaining to the freshwater media in the forms of vitamin B_12_, thiamine HCl, and biotin were omitted from the media included in this study.

**Table 1:** Buffer concentration (mM), buffering pH range, and pKa for each of the five freshwater culture media used in this study (CT, C, CSi, CB, BG-11).

| **Media** | **[Buffer] / L** | **pH Range** | **pKa at 20°C** |
| --- | --- | --- | --- |
| CT | 1.64 mM TAPS | 7.7-9.1 | 8.49 |
| C | 4.13 mM Tris | 7.0-9.0 | 8.20 |
| CSi | 2.10 mM HEPES | 6.8-8.2 | 7.55 |
| CB | 3.06 mM Bicine | 7.6-9.0 | 8.35 |
| BG-11 | 1.75mM K<sub>2</sub>HPO<sub>4</sub>•3H<sub>2</sub>O | 8.2-9.6 | 7.21 |

### Effects of flaming on media types

Twenty-five mL of each medium were aliquoted into fifty mL sterile, acid-washed glass culture tubes. For each growth medium, triplicate control and flame-treated replicates were generated. The flaming technique used in this experiment was performed in accordance with protocols expected during standard lab-based subsampling (for cell number, etc.) during a typical *M. aeruginosa* growth study. Subsampling consists of the removal of a small volume of the culture daily for cell number enumeration, though subsampling was not performed in this study. Flame replicates were inverted three times (normally done to ensure cell suspension) and held at a forty-five-degree angle while passed above the blue cone of the flame for four single events. After a brief pause, during which one would remove a small volume for cell number or inoculate cells into the media, the mouth of the culture tube was again passed through the flame four times. The cap was then immediately replaced, and the tube inverted three times before it was replaced in the rack (see <u>aseptic flaming demonstration</u> video). Controls were subject to the same procedure as denoted above, with the exception of the presence of a flame. All samples were stored at 20.5° C and approximately 15-20 μmol photons m^-2^ s^-1^. This mock sampling of uninoculated growth media was performed daily for ten days. Prior to the daily flaming, pH was measured for each replicate utilizing a pH probe, with sterilization of the probe performed prior to each sampling using 70% EtOH.

### Effects of buffer age on media pH

We assessed whether buffer age had any effect on the buffering capacity. Twenty-five mL volumes of CT media containing aged TAPS buffer stock (TAPS stock solution aged approximately three and a half months and stored at 4° C) and CT media containing fresh TAPS buffer (TAPS stock solution prepared three days prior and stored at 4° C) were aliquoted into fifty mL glass culture tubes in triplicate manner for the control treatments. The culture tubes were air-flamed in the same manner as mentioned prior. All replicates were stored on a lab bench at ∼20.5° C. The pH of replicates was monitored daily for ten days.

### Influence of gas-exchange on media pH

The potential implications of gas exchange with the atmosphere were analyzed, with twenty-five mL volumes of CT media aliquoted into fifty mL glass culture tubes in triplicate fashion for control treatments. The lids were left firmly screwed for one treatment group, while the lids were left loose for the second treatment. Replicates were stored on a lab bench at 20.5° C and 15-20 μmol photons m^-2^ s^-1^. The pH of all replicates was monitored daily for a ten-d duration. Efforts to explore potential mitigatory-practices in reducing flaming effects on media pH were performed by abstaining from any type of inversion or simulated shaking of the media. Twenty-five mL volumes of CT media were aliquoted in triplicate fashion into fifty mL glass culture tubes. Control and flamed treatments were applied daily for ten-d, without inversion of the tubes or any other form of disturbance. All replicates were stored on a lab bench at 20.5 ° C and approximately 15-20 μmol photons m^-2^ s^-1^ light intensity.

### Consequences of photo-oxidation on media pH

It has been previously demonstrated that light exposure causes photo-oxidation (and the inherent generation of reactive oxygen species such as HOOH) within a variety of media utilizing buffers such as HEPES, TAPS, Bicine, and TRIS (Morris and Zinser 2013). To determine the effects of photo-oxidation on media pH, twenty-five mL volumes of CT media were aliquoted into fifty mL glass culture tubes in triplicate form for air-flame treatments. All replicates were stored on a lab bench at 20.5° C and approximately 15-20 μmol photons m^-2^ s^-1^ light intensity. The “non-photo-oxidation” treatment replicates were wrapped in foil (kept in dark conditions) to prevent superoxide generation. The pH of all replicates was monitored daily for ten days.

### Effects of increased buffer concentration on media pH

Attempts to minimize the effects of flaming upon media pH were made by increasing the TAPS buffer concentration in CT media by ten-fold and one hundred-fold (the later serving as the positive control) and monitoring the effects of flaming and air-flaming upon the triplicated experimental groups. All replicates were stored on a lab bench at 20.5° C and 15-20 μmol photons m^-2^ s^-1^. The pH of all replicates was monitored daily for a ten-d duration.

### Testing drivers of pH decline as a result of flaming

To test the potential of flame-generated CO_2_ as the primary driver of pH decline, Limewater was prepared following standard protocol (Shakhashiri, 1983) and aliquoted into 25 mL volumes in triplicate manner for flaming in the following gradient: flamed four times, eight times, twelve times, and sixteen times, with four times meaning four passes through the flame. Negative controls served as three tubes which were air-flamed, with positive controls serving as three which were exposed to ten seconds of CO_2_ bubbling (administered *via* exhalation into straw submerged in Limewater). Qualitative analyses using turbidity due to the formation of CaCO_3_ as an indirect proxy for CO_2_ introduction were made utilizing a spectrophotometer (Thermo Spectronic Genesys 20) at 600 nm. Quantification of the net CO_2_ (TOC) incorporated into media due to flaming was performed utilizing a Shimadzu TOC analyzer. Twenty-five mL volumes of CT media were aliquoted into fifty mL acid washed, sterilized tubes in triplicate per treatment. Control, flamed four times, flamed eight times, flamed twelve times, and flamed sixteen times treatments were administered, then immediately analyzed for TOC with technical replicates of three performed per sample.

### Effects of flaming on freshwater cyanobacterial cultures

The assessment of the potential consequences flaming may have either due to H_2_O_2_ or CO_2_ on freshwater cyanobacterial cultures was performed utilizing axenic *Microcystis aeruginosa* NIES 843, which were inoculated in triplicate into CT media at a pH of 8.2 then flamed daily for ten days to mimic subsampling flaming conditions during growth assays. Note one mock-sampling event corresponds to eight total passes through the flame. Cultures were incubated at 26° C and approximately 55-60 μmol photons m^-2^ s^-1^ light intensity in incubators (VWR low temperature diurnal illumination incubator). Fluorescence was quantified daily as a proxy for cell number and cell health utilizing a fluorometer (Turner Designs TD-700). Flaming/air-flaming was performed post-fluorescence measurements. Final pH values were measured on day ten.

## Assessment

### Effects of Flaming on pH of freshwater media

After flaming, a significant decline in pH was observed in all the freshwater media analyzed in this study compared to the controls (**Figure 1).** After ten days, CT (p = 0.01) and BG-11(p = 0.02) flamed replicates experienced the greatest decline in pH, from 8.2 to average final pHs of 6.47 and 6.45, respectively **(Figure 1d, e)**. TRIS (p = 0.03), HEPES (p = 0.01), and Bicine buffered-C media (p = 0.01) had average final pH values of 7.02, 6.86, and 6.87, respectively **(Figure 1a, b, c).** Significant differences between the controls and flamed replicates manifested after only two days of daily flame sterilization (or 16 total passes through the flame). Upon the comparison of net decline in pH of each media/buffer, CT and BG-11 demonstrated similar pH declines (p = 0.99) in flame replicates, as well as control replicates (p = 0.27) **(Figure 2).** This data confirms CT (TAPS buffer) and BG-11 (inorganic phosphate buffer) demonstrated the greatest decline in pH in flamed replicates (1.5 - 1.7 pH units). TRIS, HEPES, and Bicine-buffered media replicates exhibited similar statistically significant declines in pH in control and flame replicates on a smaller scale, with no statistical difference between one another **(Figure 2).** The data demonstrated a consistent trend amongst the five-freshwater media/buffers tested, in which flaming results in a significant decline in pH after just two days and continuing up to ten days.

**Figure 1:**
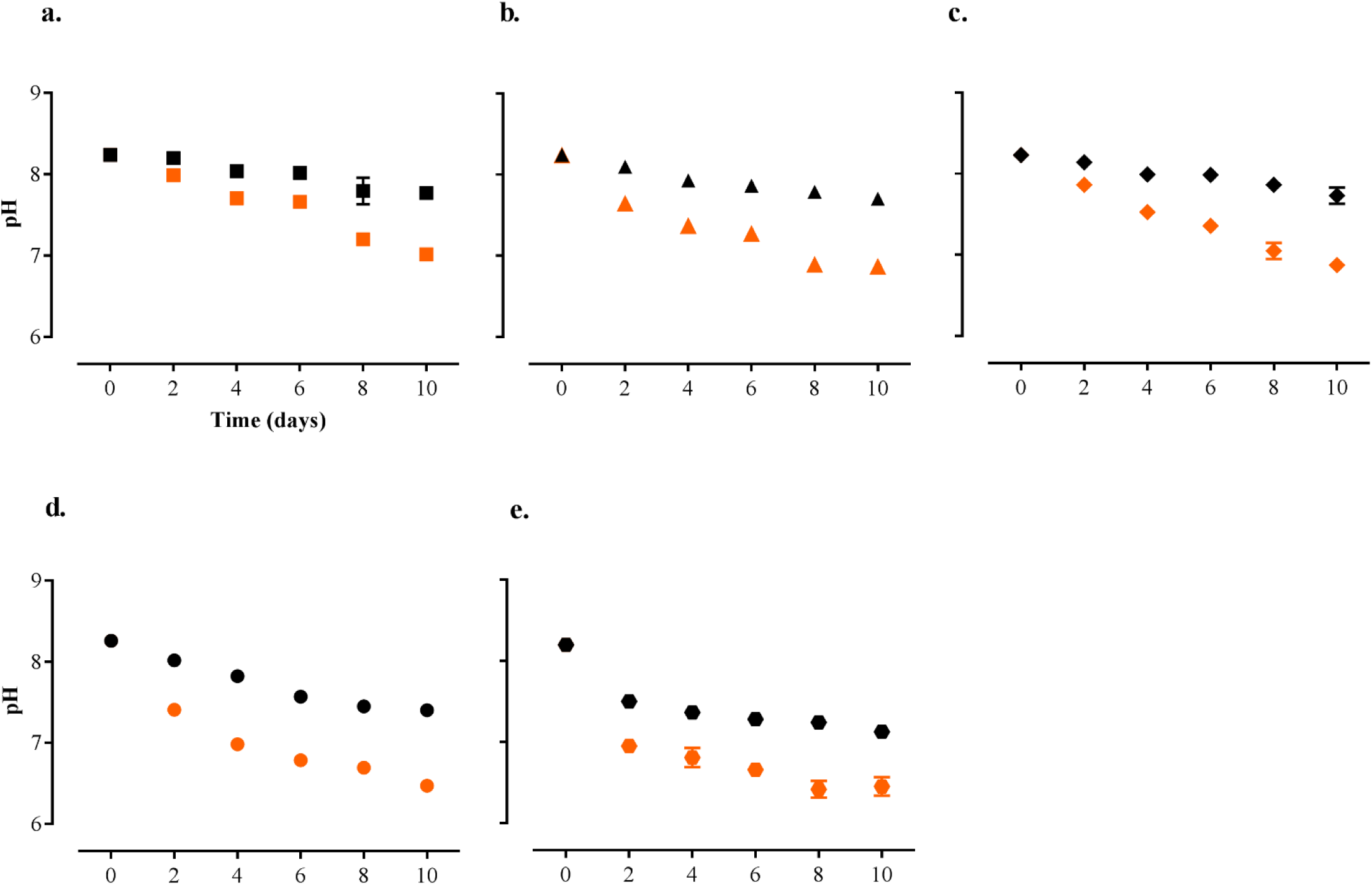
Flame-induced declines in pH in control (black) and flamed replicates (orange) of freshwater media. Data are for mean ± SEM. Where not shown, error bars are within the symbol **A.)** C media/Tris buffer (indicated by squares). **B.)** CSi/HEPES media (indicated by triangles). **C.)** CB/Bicine media (indicated by diamonds). **D.)** CT media/TAPS buffer (indicated by circle symbols). **E.)** BG-11/Inorganic phosphate buffer (indicated by hexagons).

**Figure 2:**
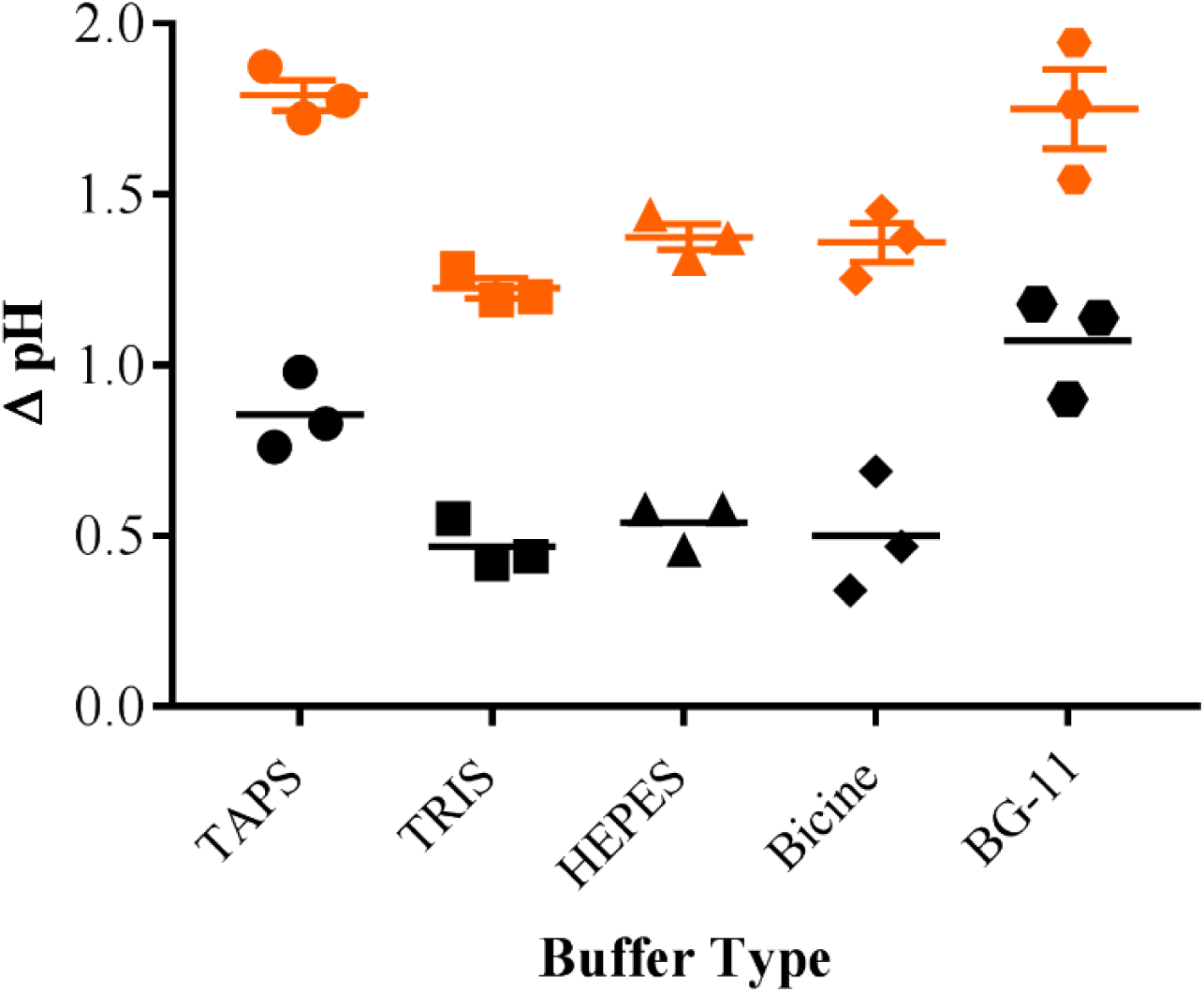
Net change in pH (T0-T10 pH) observed in each media after 10 days of aseptic flaming. Net change in pH observed in the control replicates of each media indicated in black. Net change in pH of the flamed replicates for each media indicated in orange.

Interestingly, this trend was observed (albeit to a lesser extent) when this experiment was replicated by the flaming of three media used for heterotrophic bacteria: (LB), Nutrient broth, and M9 (**Supplemental Figure 1).** In the case of the unbuffered media (LB and Nutrient broth) the pH trend was observed on a smaller scale, with flamed LB decreasing in pH from 8.2 to an average of 7.5 after ten flaming events, and Nutrient broth averaging a pH of 7.42 **(Supplemental Figure 1a, b).** M9 media demonstrated a similar decline in pH upon flaming **(Supplemental Figure 1c).** However, all time points of the M9 flamed replicates (T2-T10) were significantly different from the controls (p < 0.01) whereas this was not the case for LB and Nutrient broth. This data demonstrates flaming results in lower magnitudes of pH declines in unbuffered media in comparison to buffered media, with freshwater media experiencing the highest decline in pH due to flaming.

### Influence of environmental conditions on pH of freshwater media

Interestingly, pH decline was observed amongst control replicates in all five-freshwater media, though to a much lesser extent **(Figure 2).** Environmental factors pertaining to media handling and storage were analyzed in attempt to identify potential confounding variables/contributing factors to this pH decline observed in control replicates. Buffer age appeared to have no immediate implications corresponding to pH **(Supplemental Figure 2a).** The effects of photo-oxidation in respect to pH revealed no statistically significant difference (p = 0.99) in pH between CT replicates subjected to photo-oxidation and those exposed to photo-oxidation inhibition **(Supplemental Figure 2c).** Influences on gas exchange revealed no significant effects on pH (p = 0.90) throughout the ten-d sampling **(Supplemental Figure 2b).** Efforts to minimize pH decline in CT media were made by forming a CT -TAPS buffer concentration gradient. Daily flaming events over the course of ten days revealed the largest pH declines in the control (protocol-standard concentration of TAPS), with ten-fold and one hundred-fold increases in TAPS resulting in more stable pH **(Supplemental Figure 4a, b).** While the control TAPS concentration was not statistically different from the ten-fold increase in TAPS in both control (p > 0.07) and flamed replicates (p > 0.35), pH decline was observed at a lesser extent in the ten-fold TAPS increase replicates. Additional means of minimizing flame-induced pH decline by abstaining from tube inversion/disturbance throughout 10-d revealed a significantly lower net decline in pH in CT media **(Supplemental Figure 5).**

### Analysis of carbon dioxide as driver of flame-induced pH declines

Preliminary qualitative means for determining the absence/presence of CO_2_ in flamed media were conducted utilizing a Limewater turbidity assay (Shakhashiri, 1983). Upon flaming, indirect evidence of CO_2_ incorporation was observed via the visible formation of a calcium carbonate precipitate in the corresponding manner: negative controls or “air-flamed” replicates depicted no turbidity or precipitates, while four-times flamed, eight-times flamed, and twelve-times flamed each indicated increasing turbidities and precipitate formation, serving as a visual proxy for CO_2_ **(Supplemental Figure 3a).** To further quantify these results, spectrophotometric analyses were performed in two technical replicates of three replicates each. This data demonstrates the same trend seen in the qualitative Limewater analyses **(Supplemental Figure 3b)**, confirming the presence of CO_2._ As the number of flaming events increases, a parallel rise in turbidity and precipitate formation due to the reaction of CO_2_ with calcium hydroxide is observed. An increase in variability of CO_2_ introduction amongst replicates was observed as the number of flaming events increased, indicating CO_2_ generation due to flaming is in itself may be susceptible to variability, or the formation of the precipitate itself is variable. Quantification of the net carbon input revealed an average of 3.49 mmol C present in the controls, 4.82 mmol C present in the flamed four-times flamed replicates, 4.91 mmol C present in the eight-times flamed replicates, 5.28 mmol C present in the twelve-times flamed replicates, and 5.394 mmol C present in the sixteen-times flamed replicates **(Figure 3a).** A total of 0.79 mmol of C was found present in undisturbed CT media alone, and thus it was determined inverting the control tubes alone results in approximately 3.49 mmol C introduction. Hence, these measurements were utilized to calculate the approximate C incorporated into the replicates due solely as a result of flaming. It was determined approximately 1.33 mmol C was introduced after flamed four-times, 1.42 mmol C is introduced after flamed eight-times, 1.79 mmol C is introduced after flamed twelve-times, and 1.90 mmol is introduced after flamed sixteen-times. It was determined all flaming treatments resulted in a statistically significant (p < 0.002) increase in C incorporation compared to the controls, with increased flaming events highly correlated with a linear increase in TOC (R^2^ = 0.93, slope = 0.063) **Figure 3b).**

**Figure 3:**
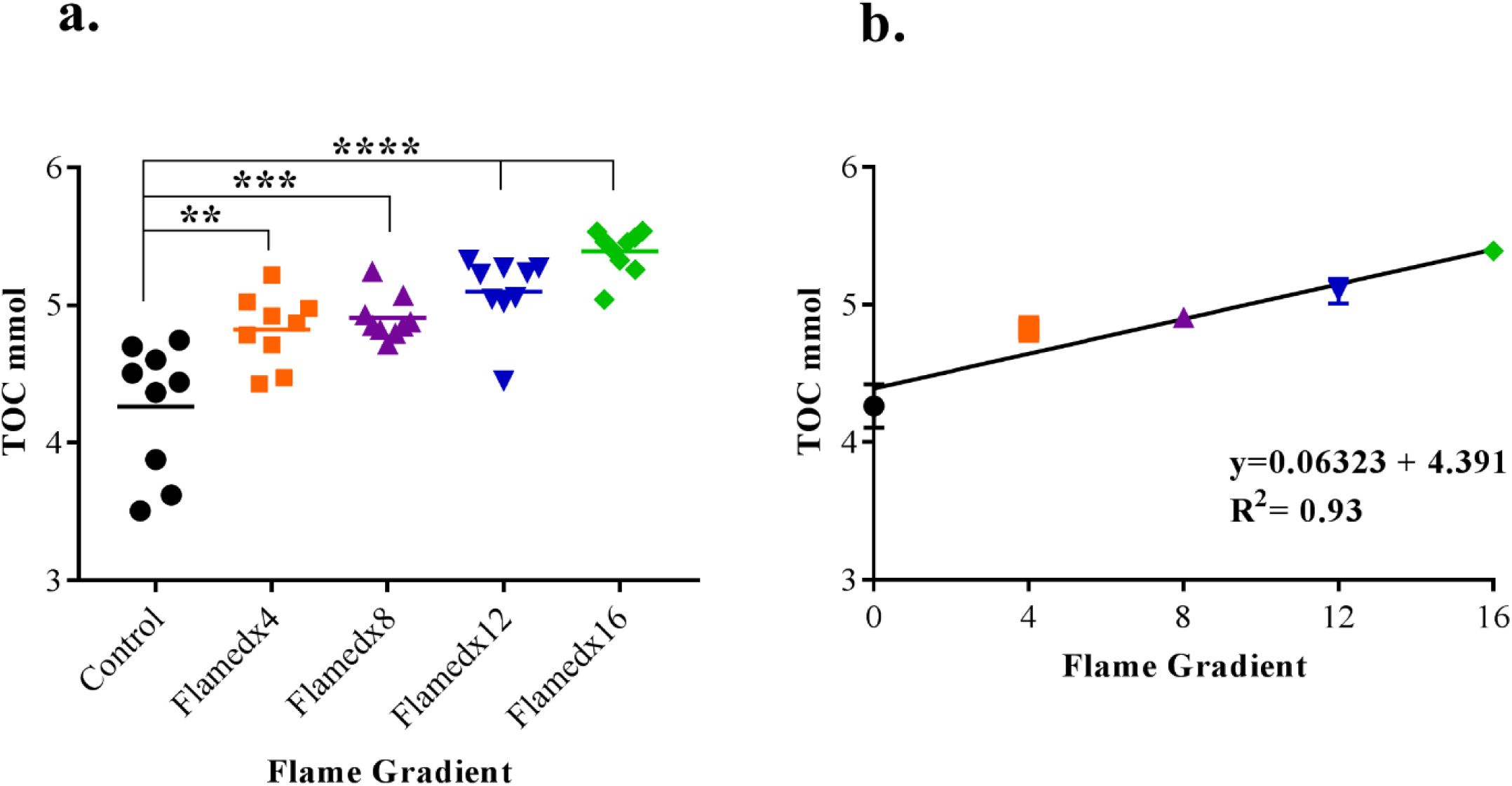
TOC (mmol) in control and flamed CT media replicates. **A.)** TOC (mmol) incorporated into CT media control and flame treatments. Flamed x4 indicated by orange squares, flamed x8 indicated by purple triangles, flamed x12 indicated by blue inverted triangles, flamed x16 indicated by green diamonds. **B.)** Linear trend line of mean TOC (mmol) introduced per flame treatment (y = 0.06323x + 4.391. R^2^ = 0.93).

### Implications of flaming for freshwater cyanobacteria cultures

Surprisingly, the control replicates of *M. aeruginosa* in CT consistently had the lowest fluorescence readings, with replicates flamed four-times and eight times exhibiting similar increased trends in fluorescence **(Figure 4a).** Unexpectedly, replicates flamed twelve-times had the highest fluorescence readings, which are indirectly indicative of cell number and biomass, while directly suggestive of cell health. Replicates that were flamed exhibited higher instances of culture death in comparison to the controls, with the replicate death observed in the flamed four-time treatment group graphed separately **(Figure 4a).** Upon the conclusion of the growth study, it was found the pH of the dead replicate was significantly lower than the other replicates which widely remained in close proximity to the initial pH of 8.2 (**Figure 4c).** Interestingly, while a replicate in the flamed eight-time treatment had a correspondingly low pH, the fluorescence of the culture did not indicate death, however it did exhibit lower fluorescence. Upon the calculation and statistical analysis of the growth rates per treatment, it was found the flamed x4 (p = 0.002) and flamed x12 replicates (p = 0.001) had a statistically higher growth rates in comparison to the control *M. aeruginosa* replicates, with replicates flamed eight-times falling short of statistical difference (p = 0.072) **(Figure 4b).**

**Figure 4:**
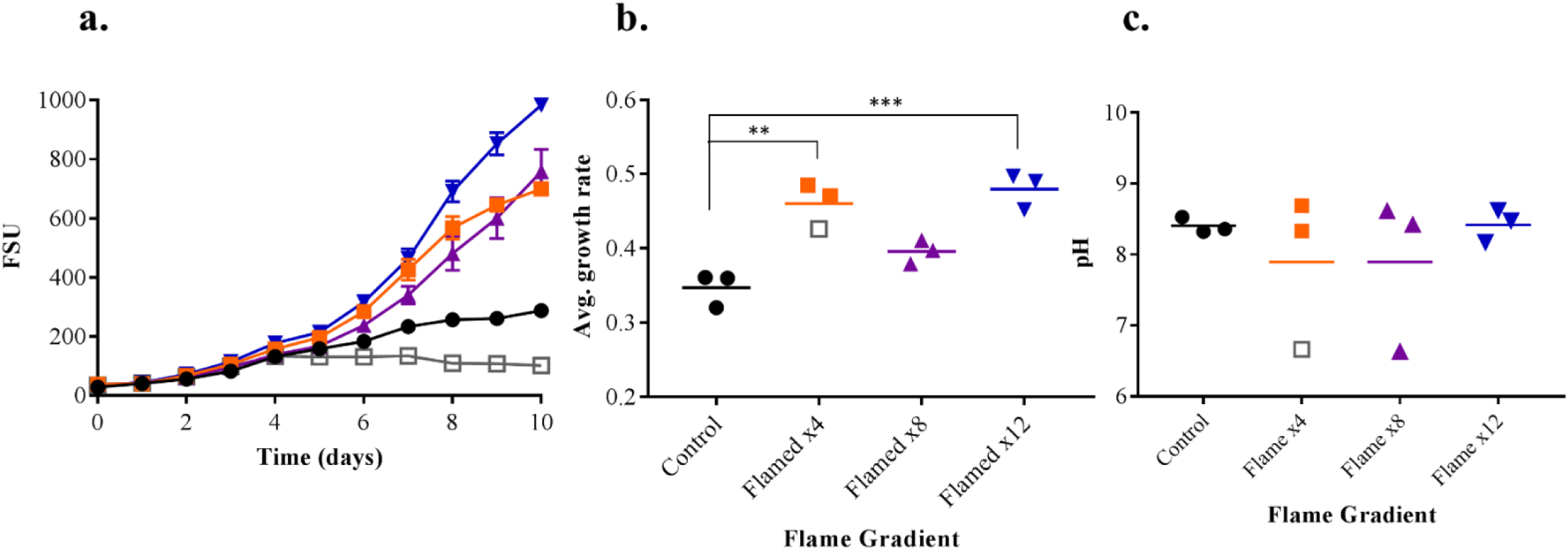
Flaming effects on CT media + *M. aeruginosa axenic culture.* **A.)** *M. aeruginosa* growth treated with flaming events: control replicates indicated by solid black dots, flamed x4 indicated by solid orange squares, flamed x8 indicated by solid purple triangles, flamed x12 indicated by solid blue inverted triangles, dead replicates indicated in grey unfilled corresponding treatment symbol. **B.)** Growth rate of each replicate + mean growth rate. **C.)** pH of replicates after 10 flaming events.

## Discussion

There is little doubt of the critical importance of aseptic technique in microbiology and microbial ecology. This is especially true when measuring microbial processes – efforts that now commonly provide rates that can drive models of global scale processes (e.g., Weitz et al. 2015). In the current study our efforts to look at microscale changes in pH as drivers of phytoplankton community composition in lakes have revealed a Heisenberg-like moment (Wheeler et al. 1983) in biology – where the process of making the measurement shapes the actual measure one is making. In the current study, our use of aseptic technique during experiments designed to look at pH effects was affecting pH. These observations are critical in face of the massive number of ongoing studies in aquatic systems where climate change associated pH is considered a major variable.

The data we have collected suggest flaming as a part of aseptic technique had substantial effects on the five freshwater culture media we tested as well as additional effects on the growth of the cyanobacterium *Microcystis aeruginosa*. While flaming is sterilization method which has been enthusiastically endorsed for biological studies for decades (Andersen 2005; Madigan 2015), our results indicate media pH was altered by as much as 1.7 units *via* the introduction of CO_2_, serving as a confounding variable. Furthermore, the assessment of experimental and environmental variables such as buffer age, photo-oxidation, and gas exchange were found to have no statistically significant effect on pH. These observations effectively ruled out additional contributors to the observed pH decline and indicated flaming was solely responsible for the observed pH declines. Previous studies have identified pH as a parameter of importance when culturing algae, with broad growth rate optima falling within pH values of 7 to 9, though many organisms exhibit growth outside of these ranges (Berberoglu et al. 2008; Huang et al. 2017; Lavens and Sorgeloos 1996). The effects of pH shifts can be extensive in algal cultures, as most algal species experience optimum growth optima at narrow and specified pH ranges (Lavens and Sorgeloos 1996; Tsaloglou 2016). The cyanobacterium used in this study, *M. aeruginosa*, has an optimal pH for growth between 7.5 - 10 (Fang et al. 2018) with *M. aeruginosa’s* optimal photosynthetic potential at pHs higher than 8.0 (Bano and Siddiqui 2004). Flaming of freshwater media, specifically during the process of aliquoting media volumes for inoculation, has the capacity to drive down the pH well below the range for optimal photosynthetic processes and growth **(Figure 1a-e).** Additionally, flaming cultures post-inoculation would further decrease pH levels if CO_2_ is not assimilated by the cyanobacteria, which may detrimentally effect culture growth and contribute to cell death. BG-11 and CT media exhibited the highest net declines in pH **(Figure 2)**, which raises significant cause for concern as these are frequently utilized freshwater media. Random culture death in certain flamed cultures **(Figure 4)** may be due to potential variability of CO_2_ incorporation from flaming, as some cultures may experience large-scale CO_2_ inputs that results in pH declines if the CO_2_ is not consumed by the culture. Additionally, previous studies have found that up to 2 μM of HOOH may be generated as a result of flaming (Morris and Zinser 2013). While it is recognized mM concentrations are needed to invoke cell death (Alam et al. 2001; Palenik et al. 1991), the 2 μM of HOOH generated from flaming may result in physiological changes from energetic draining of cells during upregulation of intracellular peroxidases (Price and Harrison 1988). This depletion of energy assets, when coupled with declining pH, serve as probable contributors to the randomized culture death and variation observed in previous growth assays. The potential contributory effect of HOOH (a weak acid) to the flame-induced pH declines observed in this research was disproven in this study, as titrations using 3% HOOH of CT to experimentally determined pHs of flamed CT required large inputs of approximately 15 mmol (data not shown).

Dissolved inorganic carbon (DIC) plays a significant role in the growth and development of phytoplankton species. Carbon dioxide and bicarbonate serve as the two major forms of carbon input in the photosynthesis reaction (Schindler et al. 1971; Wetzel 2001). Additionally, CO_2_ acts as an abiotic pH depressant, in addition to a metabolic input and product to a variety of aquatic biota (Talling 2010). Previous studies have indicated free CO_2_ is labile and readily accessible to most algae and aquatic photoautotrophs (Wetzel 2001). *M. aeruginosa* has been shown to have a high affinity for DIC which is reflective in its enzymatic low half-saturation constant, allowing it to outcompete other algal species due to its efficient carbon acquisition (Yamamoto and Nakahara 2005).

It appeared that cultures which received CO_2_ via flaming benefited substantially and exhibited higher growth rates in our study. This suggests carbon availability may be growth limiting in CT media when cultures are not bubbled or shaken **(Figure 4).** CO_2_ is extremely soluble in aqueous solutions, demonstrating two hundred-times higher solubility in juxtaposition to oxygen (Wetzel 2001). Carbonic acid in itself is a weak acid which readily and rapidly dissociates, losing its two protons in a two-step process (pKa_1_=6.43, pKa_2_=10.43 at 15° C) (Schindler 1971) (Wetzel 2001). Hence, CO_2_ that is not readily taken-up by *M. aeruginosa* in its initial gaseous form readily dissolves into the media and hydrates into carbonic acid, as demonstrated in the flaming of freshwater media alone **(Figure 1).** In addition, *M. aeruginosa’s* direct uptake of flame-generated gaseous CO_2_ is further supported in the consistent pHs amongst control and flamed replicates, as healthy cultures did not experience any significant declines in pH, save for one replicate in the flamed eight-times treatment **(Figure 4c).** Yet, when culture death did occur, the final pH was in the same range as CT media subjected to ten flaming events **(Figure 1d)**, indicating CO_2_ dissolved into the media. Should the concentration of dissolved carbon dioxide (DCD) become too high, carbonic acid formation will yield low culture pH, which would inhibit growth and development in a variety of algal species, including *M. aeruginosa* (Weisse and Stadler 2006). Thus, while CO_2_ influxes via flaming may beneficially increase growth rate and biomass in carbon-limited cultures, should the CO_2_ dissolution rate become too high, significant drops in pH will occur, implicating detrimental effects on the culture. In total it appears likely that the availability of CO_2_ coupled to pH decline shapes the inconsistency observed amongst replicates in previous experiments.

## Comments and recommendations

It has been demonstrated flaming results in a significant decline in freshwater media pH, which is attributed to the dissolution and hydration of CO_2_ into the media generating carbonic acid. This decline in pH has negative implications, as optimal pH ranges for algal photosynthesis and growth are often narrow and well-defined (Huang et al. 2018). This pH shift would also effect the bioavailability and accessibility of macro and micronutrients, including crucial trace metals within the media, many of which are associated with photosynthetic processes (Wetzel 2001). Low pH levels as a result of CO_2_ introduction, coupled with HOOH generation via flaming and excessive trace metal bioavailability, may result in randomized spontaneous cell and culture death, as seen in our previous flamed growth assays **(Figure 4).** Flaming further effects cultures by serving as a variable yet direct source of CO_2_, which is labile and thus readily accessible for uptake by *M. aeruginosa*. This CO_2_ is likely readily consumed by *M. aeruginosa* cells, increasing cell growth rate, biomass, and fluorescence **(Figure 4).** This mechanism is further supported by the relatively constant pH levels observed in all *M. aeruginosa* cultures (flamed and non-flamed) as after 10 days the pH remains constant excluding the dead replicates who demonstrate significantly lower pH’s, as lower viable cell numbers likely resulted in higher CO_2_ dissolution into the media. These CO_2_ inputs may result in a higher degree of replicate variability in unshaken *M. aeruginosa* cultures in contrast to shaken cultures, as unshaken cultures in potentially carbon-limiting media such as CT are likely to be carbon-limited. These results imply C media (in addition to CT, CB and CSi) may be carbon limiting in *M. aeruginosa* culture studies as there is no significant carbon source in the media protocol, calling for a stoichiometric re-examination of the classical freshwater media which scientists have relied upon for decades.

There is a discernable need for modern and updated aseptic alternatives to flame-induced sterilization. It is advisable to abstain from flaming when feasible, and alter flaming practices when flaming is necessary. Allowing the flamed tube to cool before replacing the cap may decrease CO_2_ incorporation into the medium. Additionally, refraining from inverting, shaking, or otherwise disturbing the media post-flaming may decrease the dissolution of CO_2_ into the media and mitigate pH decline, though this is not always feasible. When CT media was subject to a ten-d flaming study without any inversion/shaking, it was found the pH decline was significantly diminished **(Supplemental Figure 5)** to resemble pH declines not significantly different (p>0.30) from the non-flamed controls. Additionally, exercising caution in media selections is advised, as the commonly utilized medium (BG-11) was shown to have the highest decline in pH overall. While this medium’s primary buffer, potassium phosphate, is said to be an exceedingly strong buffer with a high buffering capacity (Sigma-Aldrich), this appears to be contradicted in this study. Hence, this medium should be avoided, or adapted to mitigate the dramatic pH declines as demonstrated in this study **(Figure 1e).** Ten-fold increases in TAPS buffer concentration within CT media have been shown to mitigate pH shifts within un-flamed replicates **(Supplemental Figure 4)** with no biological consequences **(Supplemental Figure 6).** However, the application of this approach to varying media must be pursued with caution, as TRIS is toxic to cells in high concentrations and HEPES generates higher amounts of reactive oxygen species when present in higher concentrations (Morris and Zinser 2013). Potassium phosphate dibasic trihydrate has also been shown to negatively affect ligands, proteins and restriction enzyme activity while leading to the precipitation of calcium and magnesium (Sigma Aldrich, 2019). The Bis(2-hydroxyethyl) amine family of buffers has been found to complex with a variety of metals common in biological and environmental studies, resulting in advisories against use in biological studies (Ferreira et al. 2015). Hence, our results indicate a ten-fold buffer increase in CT may serve as the most suitable media for freshwater culture work, specifically in future studies concerning pH or carbon such as climate change research.

This study serves as a cautionary of the unintended effects of flaming of culture vessel openings upon the media and microorganisms. While flaming has served as an aseptic technique for decades, it may be time to put aside the Bunsen burner and pursue further alternatives to this classic practice.

## Supporting information

Supplementary information for Zepernick et al. 2019

## Acknowledgements

We thank Dr Gary LeCleir, Eric Gann, Robbie Martin, Lena Pound, Naomi Gilbert and Professor George Bullerjahn for comments. This work was made possible due to grants from the National Science Foundation (DEB-1240870; IOS-1451528) to SWW. This work was also supported by funding from the NIH (1P01ES028939-01) and NSF (OCE-1840715) to the Bowling Green State University *Great Lakes Center for Fresh Waters and Human Health*.

