## Supplementary information for Zepernick et al. 2019 for "Flaming as part of aseptic technique increases CO_2 (g)_ and decreases pH in freshwater culture media"

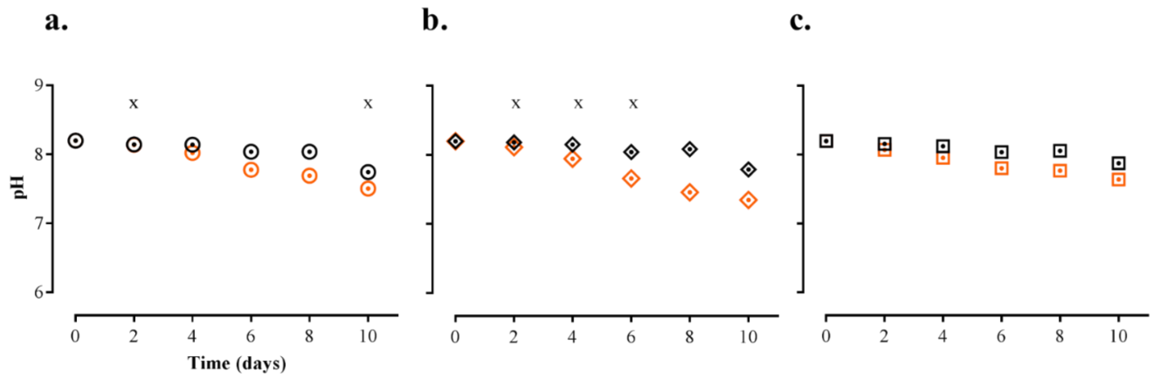


**Supplemental Figure 1.** Effects of flaming on heterotrophic media **A.)** Effects on pH of LB media, control indicated by black circles, flaming indicated by orange circles **B.)** Effects on pH of Nutrient broth, control replicates indicated by black triangles, flamed replicates indicated by orange triangles **C.)** Effects on pH of M9, control replicates indicated by black squares, flamed replicates indicated by orange squares. Control and flamed replicate pHs that demonstrate no statistical difference denoted by x's.

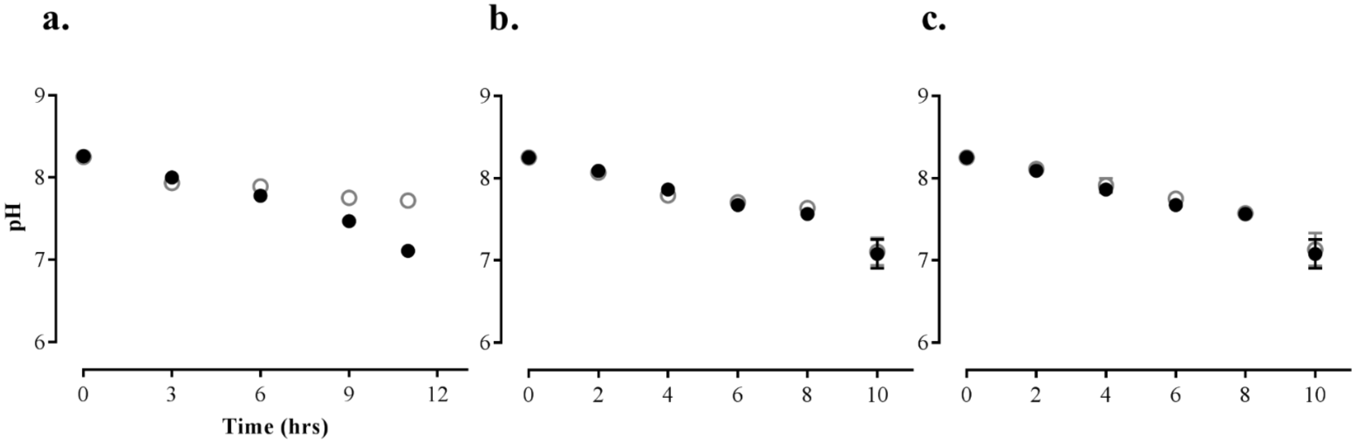


**Supplemental Figure 2:** Effects of environmental conditions on pH of CT media. **A.)** Effects of buffer age on pH, aged buffer indicated by solid black circles, new buffer indicated by open grey circles **B.)** Effects of gas exchange on pH, closed caps indicated by solid black circles, loosened caps indicated by open grey circles **C.)** Effects of photo-oxidation on pH, light-exposed samples indicated with solid black circles, dark-treatment samples indicated with open grey circles.

**a.**

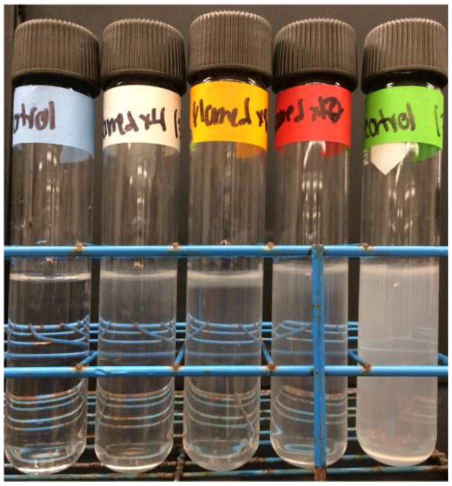

**b.**

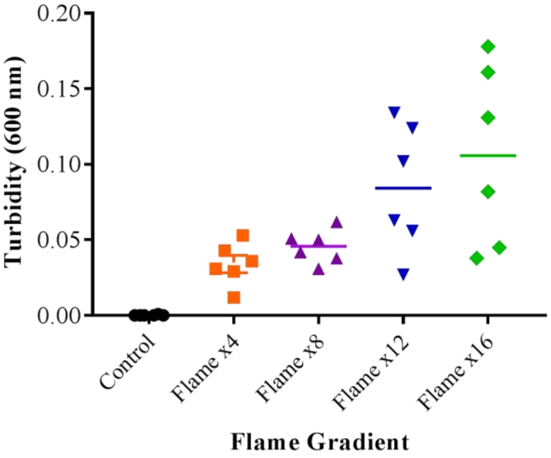


**Supplemental Figure 3:** Determination of $CO_2$ as the primary driver of pH decline in freshwater media. **A.)** Limewater turbidity assay demonstrating increased turbidity with increased flaming. Treatments from left to right:  control, flamed x4, flamed x8, flamed x12, and positive control. **B.)** Turbidity (600nm) of $CaCO_3$ precipitation as indirect proxy for $CO_2$ incorporation.

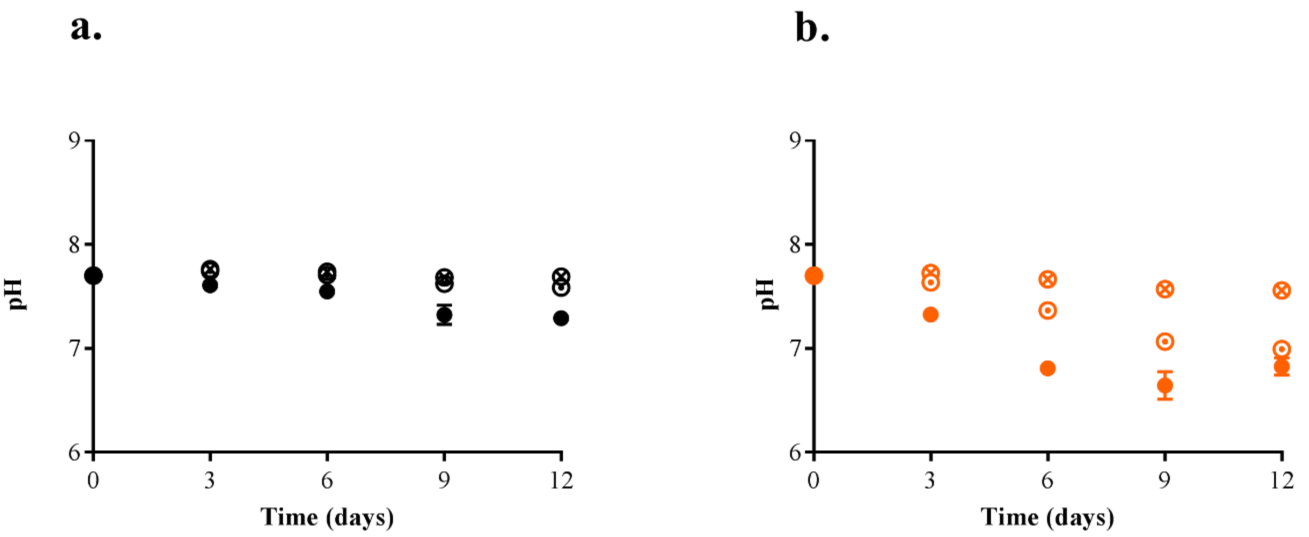


**Supplemental Figure 4:** Mitigation of pH decline utilizing CT media TAPS buffer gradient. **A.)** Effects of increased [buffer] on pH in control replicates, standard TAPS concentration indicated by solid black circles, TAPsx10 indicated by open dotted circles, TAPSx100 indicated by open crossed circles **B.)** Effects of increased [buffer] on pH in flamed replicates, with standard TAPS concentration indicated by solid orange circles, TAPsx10 indicated by open dotted circles, TAPSx100 indicated by open crossed circles.

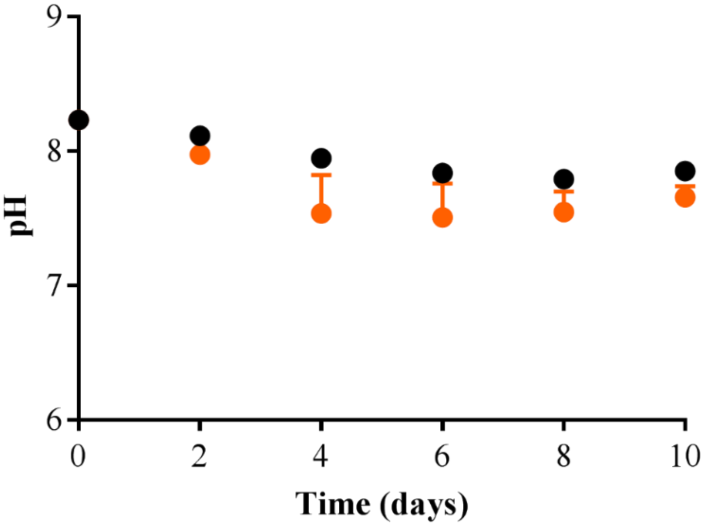


**Supplemental Figure 5:** Effects of non-inversion on control and flame replicate pH decline. Non-inverted CT media control replicates indicated by solid black circles, with non-inverted flamed CT media replicates indicated by solid orange circles.

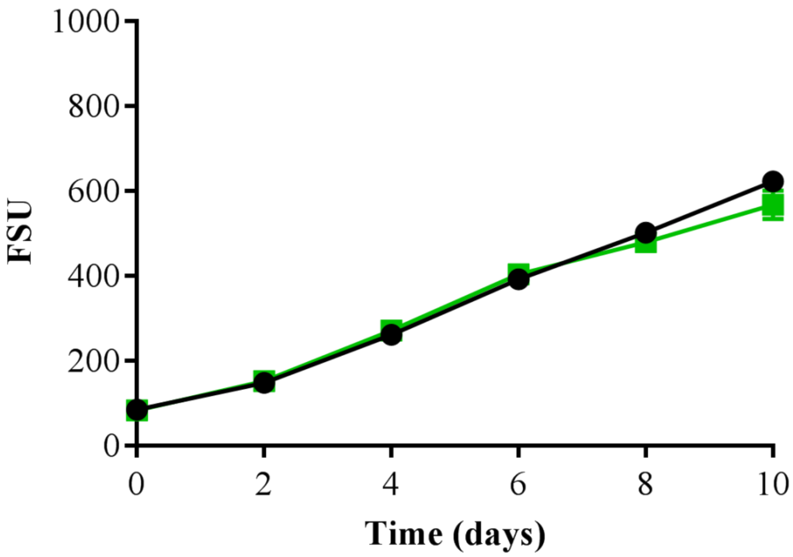


**Supplemental Figure 6:** Comparison of *M. aeruginosa* biomass accumulation and cell health (FSU) when grown in CT media with standard TAPS concentration indicated by black circles, to CT media with a ten-fold increase in TAPS concentration indicated by green squares.
